# Non-invasive characterization of human bone marrow by cell free messenger-RNA reveals response to growth factor stimulation and hematopoietic reconstitution after transplantation

**DOI:** 10.1101/516666

**Authors:** Arkaitz Ibarra, Yue Zhao, Neeraj S. Salathia, Jiali Zhuang, Vera Huang, Alexander D. Acosta, Jonathan Aballi, Shusuke Toden, Amy P. Karns, Intan Purnajo, Julianna R. Parks, Lucy Guo, James Mason, Darren Sigal, Tina S. Nova, Stephen R. Quake, Michael Nerenberg

**Affiliations:** Molecular Stethoscope, Inc., San Diego, CA 92121.; Scripps Clinic Medical Group, Scripps Green Hospital, La Jolla, CA 92037.; Department of Bioengineering and Department of Applied Physics; Stanford University and Chan Zuckerberg Biohub, Stanford, CA 94305.

## Abstract

Circulating cell free mRNA (cf-mRNA) holds great promise as a non-invasive diagnostic biomarker. However, the biological origin of cf-mRNA is still not well understood, limiting the clinical applications of this technology. Here, we use the bone marrow (BM) and pharmacologic manipulation of its resident cells as a window to study the origin of cf-mRNA. Using NGS-based profiling, we show that cf-mRNA is enriched in transcripts derived from the BM compared to circulating cells. Further, BM ablation experiments followed by hematopoietic stem cell transplants in cancer patients show that cf-mRNA levels reflect the transcriptional activity of BM resident hematopoietic lineages during marrow reconstitution. Finally, by stimulating specific BM cell populations in vivo using growth factor therapeutics (i.e. EPO, G-CSF), we show that cf-mRNA reveals dynamic functional changes in growing cell types, suggesting that, unlike other cell-free nucleic acids, cf-mRNA is secreted from living cells, rather than exclusively from apoptotic cells. Our results shed new light on the biology of cf-mRNA and demonstrate its potential applications in clinical practice.

## Main text

### Introduction

Blood is a liquid connective tissue that irrigates all organs, supplying oxygen and nutrients to the cells of the body while collecting their waste, including lipids, proteins and nucleic acids. It is well known that these circulating biomolecules contain information linked to specific organ health. While most effort has focused on circulating proteins and lipids, circulating cell free nucleic acids (cf-NA) have recently emerged as a non-invasive tool for diagnosis and monitoring of health and disease ^1^. For example, cell free DNA (cfDNA), the most well-characterized cf-NA, has been utilized for prenatal diagnostics, transplant rejection and monitoring of cancer ^2-6^. Despite these advances, the value of cf-DNA tests is mainly restricted to physiologic and disease situations characterized by genetic differences (i.e. pregnancy, transplants or tumors). In contrast, cell free mRNA (cf-mRNA) transcriptome can be considered as a compendium of transcripts collected from all organs ^7^. Since some of these circulating transcripts correspond to well-characterized tissue-specific genes, they can be used to monitor the health or state of individual tissues of origin. Indeed, cf-mRNA has been shown to reflect fetal development and predict preterm delivery in pregnant women ^7-9^ and as a cancer biomarker ^10,11^.

Strikingly, the biological processes underlying the presence of mRNA transcripts in circulation remain largely unknown. In the case of cf-DNA, studies have shown the mechanism is passive release into circulation upon cell death ^12,13^. In contrast, it is known that RNA molecules can be actively secreted from cells ^11,14-16^. Much work has focused on the secretion of smaller RNA molecules into exosomes and other lipid vesicles, and it has been shown that on a per molecule basis mRNA comprises a minor fraction of this phenomenon ^17^. It has therefore remained unresolved whether circulating cf-mRNA is primarily released from living cells or only from those cells undergoing cell death.

In this study, we conducted NGS-based whole-transcriptomic profiling of cf-mRNA and compared expression levels to those from circulating cells of the blood (CC) to decipher the origin of circulating transcripts and better understand their clinical utility. We found that most cf-mRNA transcripts are of hematopoietic origin, and demonstrate in healthy subjects and multiple myeloma patients, that cf-mRNA is enriched in bone marrow (BM) -specific transcripts. Longitudinal studies of cancer patients undergoing BM ablation and transplantation showed that cf-mRNA profiling non-invasively captures temporal transcriptional activity of the BM. Mechanistically, stimulation of specific BM-lineages with growth factor therapeutics indicates that cf-mRNA fluctuations reflects active lineage-specific transcriptional activity. Collectively, our data provide new insights into the biological origins of cf-mRNA, strongly suggesting that living cells secrete cf-mRNA, and demonstrate the potential clinical use of circulating transcripts as non-invasive biomarkers to replace BM biopsies.

## Materials and methods

### Patients

**Multiple myeloma patients** eligible for autologous marrow transplantation were recruited from the Scripps bone marrow transplant center. Patients with non-secretory disease or plasma cell leukemia were excluded. Three total patients were enrolled with daily blood draws collected throughout the cytoreductive conditioning regiment and subsequent hospital stay. High-dose melphalan was used to ablate the marrow over a 2-day conditioning regiment, followed by transplantation of hematopoietic stem cells. Sequential daily collections discontinued the day of hospital discharge. Follow-up bone marrow biopsy occurred between 60-90 days. Complete blood counts (CBCs) were collected as part of the study. Plasma was processed within 2-hours of blood collection and stored.

**Erythropoietin (EPO) treated patients** were recruited for study enrollment provided they were administered erythropoietin as part of routine medical care. Potential patients were excluded if they were 1) currently on any anti-cancer therapy; 2) had active hemolysis from any cause or 3) were pregnant. Patients were consented and enrolled from the Renal and Hematology/Oncology Clinics at Scripps Clinic Cancer Center. Per standard clinical care, a single dose of erythropoietin was administered per month. Blood was collected at day 0 (before administration of EPO), and at days 1, 4 and 10 after administration of EPO. Day 4 and day 10 collections could be adjusted +/-1 day to accommodate patients’ schedules. A subset of patients consented to an expanded protocol allowing for blood collections up to day 30. CBCs were performed as well. Cell-free hemoglobin protein (ARUP labs) and albumin levels (ARUP labs) were determined at each time point.

#### Healthy controls

Whole blood from healthy controls was obtained from the San Diego Blood Bank. Plasma/ serumwas processed within 2-hours of blood collection, frozen and stored for batch processing.

#### G-CSF Cohort

Normal healthy individuals preparing to donate peripherally harvested stem cells for allo-transplants, were recruited from Scripps and enrolled as part of our G-CSF cohort. In total, three patients were consented and donated blood during their stem cell mobilization. Two tubes of blood were collected at day 0 (before administration of G-CSF), and at days 1, 4 and 10 after administration of G-CSF. Day 4 and day 10 collections could be adjusted +/-1 day to accommodate patients’ schedules and additionally, the Day 10 collection was optional. Peripheral harvest of stem cells occurred on day 4 by leukapheresis. CBCs were performed for each sample.

#### AML Cohort

Patients with known acute myeloid leukemia (AML), in preparation for submyeloablative treatment and allogeneic stem cell transplantation as part of standard care, were recruited for daily blood draws throughout their treatment and stem cell transplant Three patients were enrolled in our study, and submyeloablative treatment were generally 6-days, using a combination of fludarabine and melphalan to obtain a partial ablation of the marrow, prior to transplantation. Hematopoietic stem cells obtained from a single donor, were administered on day 0, and daily blood draws were continued through the hospital stay. In-hospital collections were limited to day 45 post-transplant. Follow-up routine bone marrow biopsies were performed. CBCs were collected as part of standard care and the data were included in the study. Plasma was processed within 2-hours of blood collection and stored for batch processing. Two of the AML patients were monitored for ~8 weeks, while blood samples for the third patient collected until 15-day post-transplant when the patient was discharged from the hospital.

### Patient Consent

All studies were approved by their respective institutional IRBs and patients consented according to submitted study protocols. Molecular Stethoscope maintained approval for blood collection and research through Western IRB Protocol #20162748, under which normal control samples were collected. In collaboration with the Scripps Cancer Center and the Blood & Marrow Transplant Program at Scripps Green Hospital, G-CSF and EPO studies were conducted under Scripps Institutional Review Board approved protocol IRB-16-6808. Our studies involving hematopoietic bone marrow transplants, for both Multiple Myeloma and Acute Myeloid Leukemia, were approved by and conducted in accordance with Scripps IRB Protocol IRB-17-6953, in collaboration with the same groups.

### Sample processing

Blood samples were collected in EDTA tubes (BD #366643) for plasma processing or in BD Vacutainer red-top clotting tubes (BD #367820) for serum processing. cfRNA was isolated from plasma/serum, converted to cDNA, and NGS libraries were generated for Illumina sequencing. Libraries were quantified in a Quantifluor (Agilent Quantus Fluorometer, Promega) using QuantiFluor ONE dsDNA kit (Promega) and library size was checked on the Bioanalyzer (Agilent Technologies) using high sensitivity DNA chips (Agilent Technologies). Samples were pooled and sequenced on a NextSeq 500 (Illumina) platform according to manufacturer’s instructions.

### Sequence data processing, alignment and transcriptome quantification

Base-calling was performed on an Illumina BaseSpace platform, using the FASTQ Generation Application. Adaptor sequences are removed and low quality bases trimmed, using cutadapt (v1.11). Reads shorter than 15 base-pairs were excluded from subsequent analysis. Read sequences are then aligned to the human reference genome GRCh38 using STAR (v2.5.2b) with GENCODE version 24 gene models. Duplicated reads are removed by invoking the samtools (v1.3.1) rmdup command. Gene expression levels were inferred from de-duplicated BAM files using RSEM (v1.3.0).

### Differential Expression Analysis

Differential expression analysis between different conditions was performed using DESeq2 (v1.12.4). RSEM-estimated read counts are used as input for DESeq2. Genes with fewer than 20 reads across the samples are excluded from this analysis. Potential Gene Ontology enrichment of differentially expressed genes were examined using the R package limma (v3.28.21).

### Tissue/cell-type specific genes

Tissue (cell-type) specific genes are defined as genes that show much higher expression in a particular tissue (cell-type) compared to other tissues (cell-types). Information about tissue (cell-type) transcriptome expression levels was obtained from the following two public databases: GTEx (https://www.gtexportal.org/home/) for gene expression across 51 human tissues and Blueprint Epigenome (http://www.blueprint-epigenome.eu/) for gene expression across 56 human hematopoietic cell types. For each gene, the tissues (cell-types) were ranked by their expression of that particular gene and if the expression in the top tissue (cell-type) is > 20 fold higher than all the other tissues (cell-types) the gene was considered specific to the top tissue (cell-type).

### Immunoglobulin gene repertoire in multiple myeloma patients

For clone-type assembly, we performed *de novo* transcriptome assembly using Trinity. Next the assembled contigs were compared to immunoglobulin gene annotation database IMGT (http://www.imgt.org/) using igBLAST (v2.5.1) to identify the V(D)J combinations. To quantify the relative abundance of variable region genes, we collected reads that were either unaligned to the human reference genome or aligned to an annotated Ig gene by STAR and d them to the sequences in the IMGT database using igBLAST. Relative abundance was calculated as the ratio of number of reads mapped to a particular Ig gene over the total number of reads mapped to any Ig gene.

### Unsupervised clustering of Multiple Myeloma and AML samples

Genes that met the following two criteria were selected for clustering: 1) the maximum expression across time points higher than 50 TPM (transcripts per million); 2) the ratio of the highest expression over the lowest was greater than 5. For each of the selected genes, the expression values were normalized by dividing each value by the maximum value across all time points. The purpose of this normalization was to bring all the genes to a comparable scale and focus on their relative changes across time points instead of their absolute expression levels. K-means and hierarchical clustering were then performed to find genes that share similar temporal expression patterns.

### Decomposing data with non-negative matrix factorization (NMF)

Genes whose expression was lower than 20 TPM in all samples were excluded from the decomposition analysis. For each of the remaining genes, the expression values were normalized by dividing each value by the maximum value across all samples. The purpose of this normalization step is to bring all the genes to a comparable scale. NMF is then performed on the normalized values to decompose the genes into 8-12 components. NMF decomposition is implemented by invoking the “decomposition.NMF” class in the sciki-learn Python library. NMF decomposition creates groups of genes (components) sharing similar expression patterns (correlated across samples) in an un-supervised manner, thereby revealing underlying structures within the data. In order to better understand the discovered components, we select genes enriched in that particular component and examined 1) their expression levels across 51 human tissues in GTEx; 2) their expression levels across 55 human hematopoietic cell types from the Blueprint Epigenome consortium; 3) their potential Gene Ontology functional enrichment. By integrating those three sources of information we are able ascertain the tissue/cell-type origin of most components.

## Results

### cf-mRNA transcriptome is enriched in hematopoietic progenitor transcripts

To characterize the landscape of the human cell free RNA transcriptome we isolated and sequenced cf-mRNA from serum of 24 healthy donors. Among this cohort, we identified 10,357 transcripts with >1TPM (transcripts per million) and 7,386 transcripts with >5TPM in at least 80% of the samples, reflecting the high diversity and consistency of cf-mRNA transcriptome among healthy subjects (Table S1). We used non-negative matrix factorization to decompose the cf-mRNA transcriptome in an unsupervised manner ^18,19^ and gene expression reference databases (GTEx and Blueprint) to estimate the relative contributions of the different tissues and cell types (see Material and Methods). The majority of the transcripts detected in cf-mRNA, ~85% on average, are of hematopoietic origin, with the remaining ~15% being of non-hematopoietic origin (Fig 1A). Specifically, 29.7% of transcripts are of megakaryocyte/platelet origin, 27.9% are of lymphocyte origin, 12.8 % of granulocyte origin, 3% of neutrophil progenitor origin, 11.3% of erythrocyte origin and 15.2% derived from solid tissues. (Fig 1A). To gain insights into the origin of these transcripts, similar deconvolution analysis was performed in whole blood samples from 19 healthy individuals from previously reported RNA-Seq data ^20^. As expected, the whole blood transcriptome is largely composed of lymphocyte (69.1%) and granulocyte (21.8%) transcripts, with an additional 7.5% of transcripts of erythrocyte origin and minor contributions from other cell types and tissues (Fig 1A). These analyses demonstrate the higher diversity of cf-mRNA transcriptome, which contains a larger fraction of non-hematopoietic transcripts and of hematopoietic progenitor genes derived from the BM.

**Figure 1.**
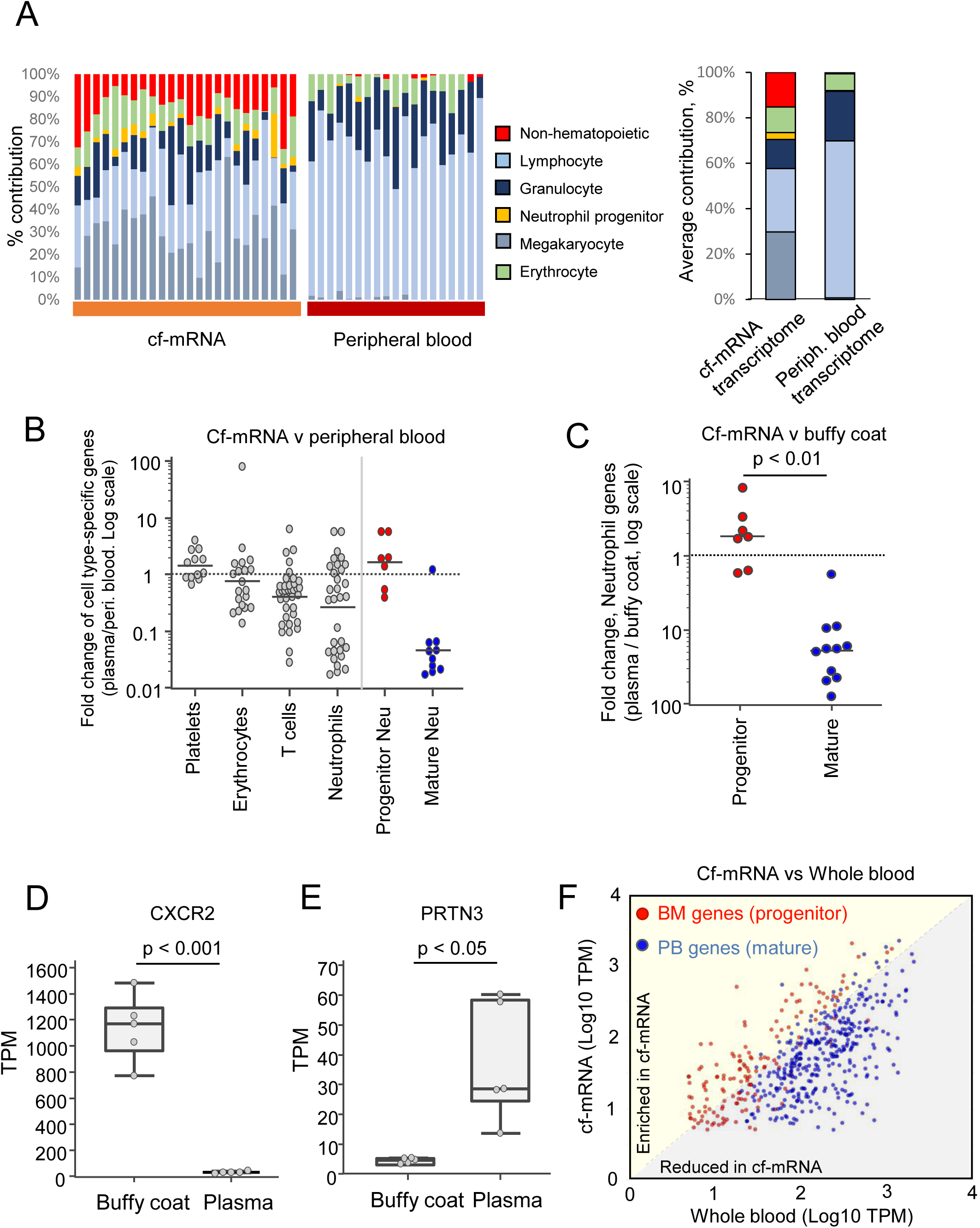
cf-mRNA transcriptome is enriched in immature hematopoietic transcripts from the bone marrow compared to circulating blood cells. A. Left panels, cf-mRNA transcriptome and whole blood transcriptome from healthy subjects was decomposed using non-negative matrix factorization and tissue contribution estimated using public databases. cf-mRNA was sequenced from 24 normal donors and whole blood RNA-Seq data from 19 healthy individuals was obtained from Nguyen et al, 2017. Estimated contribution of the indicated cell types/tissues for each sample is shown. Right panel, average values for each bio fluid (24 cf-mRNA and 19 whole blood samples) are shown using the same color code. B. RNA-seq was performed in 3 paired plasma and peripheral blood samples from healthy individuals. Levels of indicated cell type-specific transcripts were compared between cf-mRNA and peripheral blood for all 3 donors. Average fold change (cf-mRNA/peripheral blood) among the 3 individuals is represented (log scale). Red dots, neutrophil progenitor transcripts. Blue dots, mature neutrophil transcripts. Cell type specific genes were identified as explained in Materials and Methods. See also Table S2. C. RNA-seq was performed in 5 paired plasma and buffy coat samples from healthy individuals. Levels of mature and progenitor neutrophil transcripts in plasma and matching buffy coat specimens were compared. Average fold change of these transcripts (plasma/buffy coat) in the five paired samples is shown (log scale). P-value, t-test. D-E. Box-plot comparing the normalized levels (TPM) of the indicated transcripts in paired buffy coat and cf-mRNA samples measured by RNA-Seq (n=5, p-value: t-test), showing that cf-mRNA is enriched in immature (PRTN3) hematopoietic transcripts (E) and depleted of mature transcripts (CXCR2, D). F. Scatter plot comparing the levels in matching cf-mRNA (Y axis) and whole blood (X axis) of BM-specific genes (red dots) and peripheral blood-specific genes (blue dots), which form two distinct populations (p<0.001). (See also Fig S1).

To confirm the presence of BM-specific transcripts in circulation we performed RNA-Seq of 3 paired peripheral blood and plasma samples from healthy donors (Fig S1A) and compared the levels of the main hematopoietic cell type-specific transcripts (i.e. neutrophils, erythrocytes, platelets /megakaryocyte, T cells) in these specimens (Fig 1B, FigS1B-C). Striking differences were observed among neutrophil-specific transcripts (Fig1B). Using the hematopoiesis transcriptomic reference database (Blueprint), we observed that transcripts expressed in mature circulating neutrophils exhibit extremely low levels in plasma compared to peripheral blood (Fig1B). In contrast, transcripts expressed in BM-residing neutrophil progenitors are highly enriched in cf-mRNA (Fig 1B). To confirm these findings, we performed RNA-Seq of five paired plasma and buffy coat samples. Consistently, neutrophil mature and progenitor transcripts were found to form distinct populations (Fig 1C), in which cf-mRNA is depleted of mature transcripts such as the chemokine receptors CXCR1 and CXCR2 (Figure 1D, p<0.01) compared to buffy coat, but enriched in progenitor transcripts such as PRTN3 (myeloblastin precursor), CTSG (cathepsin G) and AZU1 (azurocidin precursor) (p<0.05, Fig 1E, FigS1D and FigS1E). These data support the presence of BM transcripts in cf-mRNA; indeed, quadratic programing deconvolution analysis of hematopoietic transcripts from healthy donors indicated that BM transcripts contribute ~9% of cf-mRNA transcriptome, in contrast to ~1% in whole blood.

To further confirm this result, we performed RNA-seq of BM of human origin and compared it with the peripheral blood transcriptome. We identified 377 genes enriched in BM transcriptome (>5 fold, “BM genes”), representing hematopoietic progenitors (i.e. neutrophil progenitors and mesenchymal stem cells from the BM). Interestingly, progenitor transcripts such as PRTN3, CTSG and AZU1 are among the top transcripts enriched in BM transcriptome. In addition, 374 genes were identified enriched in peripheral blood (>5 fold, “PB genes”), representing mature circulating blood cell genes, as expected (i.e. associated with mature granulocytes and lymphocytes). Subsequently, the levels of “BM genes” and “PB genes” were compared in 3 matching whole blood and plasma samples, which confirmed that these transcripts segregate into two populations (p<0.001), with cf-mRNA being enriched in hematopoietic progenitor genes (“BM genes”) and depleted of mature genes (“PB genes”) compared to whole blood (Fig 1F and FigS1F). In summary, our data indicate that the circulating transcriptome includes information quite distinct from what is found in the CC transcriptome, and confirms that cf-mRNA captures transcriptional information derived from the BM.

### Non-invasive measurement of bone marrow-specific transcripts by cf-mRNA profiling in Multiple Myeloma patients

As further evidence that BM-specific transcripts can be detected in cf-mRNA and to demonstrate their potential clinical utility, we recruited three Multiple Myeloma (MM) patients. MM is characterized by the clonal expansion and accumulation of malignant plasma cells almost exclusively in the BM. These cells express specific immunoglobulin (Ig) rearrangements, in contrast to plasma cells of healthy individuals, which express multiple Ig combinations. In this study, MM patients underwent melphalan-mediated BM ablation (starting at day-2) followed by autologous hematopoietic stem cell (HSC) infusion (day 0) (Figure 2B). We isolated and sequenced cf-mRNA from plasma of these patients before BM ablation (day −2). Clonal expansion of Ig heavy (IgH) and Ig light (IgL) chains transcripts was identified for 2 of 3 patients. For instance, in Patient 2 we detected IGHG1 and IGKC transcripts as the most prevalent Ig constant regions (Fig S2A-C). For the variable regions, Ighv3-15 and Igkv2-24 transcripts dominated the sample’s transcriptome, while no clonal lambda regions were detected (Fig2A, C, Fig S2C). In contrast, and as expected, no clonal transcripts were observed in plasma of healthy individuals (Fig 2A). Similar analyses in Patient 1 revealed a clone composed of the IgH constant chain IGHA1 and variable region IGHV1-69, and IgL lambda chain IGL1 and variable region IGLV1-40 (Fig S2D). In both cases the malignant clones we identified are consistent with the molecular testing performed from BM aspirates (Table S3). For Patient 3 we detected no dominant Ig rearrangements (data not shown), likely due to the low number of plasma cells in the BM of this Patient at the start of this study (Table S3). Malignant plasma cells are rarely found in circulation in MM patients; indeed, RNA-Seq analysis of the matching buffy coat of Patient 2 samples before chemotherapy treatment showed only very low levels of a repertoire of IgH and IgL transcripts, with no dominant rearrangements (Fig 2A, C, and Fig S2A-C), highlighting the unique ability of cf-mRNA to capture the clonal Ig transcripts generated by plasma cells in the BM.

**Figure 2.**
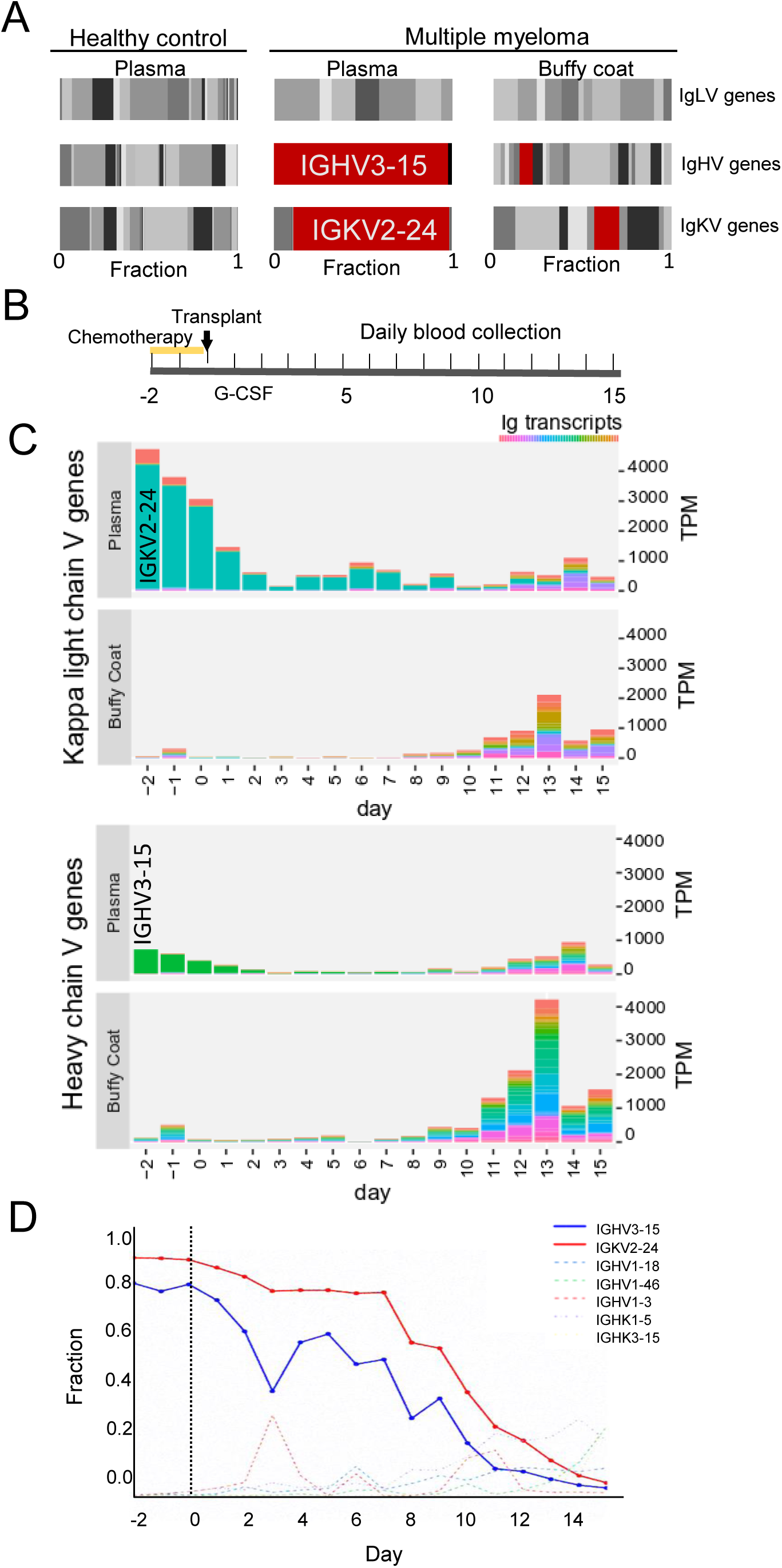
Cf-mRNA transcriptome captures Ig transcripts derived from the BM of Multiple Myeloma patients. A. Matching cf-mRNA and buffy coat samples from a Multiple Myeloma patient before BM ablation (day-2) were analyzed by RNA-Seq. Fraction of transcripts from the variable regions of the immunoglobulin heavy and light chains identified in plasma and buffy coat samples are shown (center and right panels). Clonally amplified transcripts are indicated in red and dominated the cf-mRNA of the MM Patient. Levels of Ig transcripts in plasma of a healthy individual (left panel) are shown as reference. B. Schematic of the therapeutic treatment performed in MM patients. Melphalan-mediated BM ablation started at day −2, autologous stem cell transplant was performed at day 0. Steroids and G-CSF were then administered as supportive care. Blood was collected every day during the study. C. Bar graphs showing the normalized values (TPM, Y axis) of Ig transcripts detected by RNA-Seq in paired plasma and buffy coat samples throughout the treatment. The repertoire of variable regions of Ig heavy chain and Ig Kappa light chain are shown in a color gradient. Dominant transcripts identified in plasma are indicated. Day of blood collection with respect to transplant is indicated in the X axis. D. Fraction of transcripts from variable Ig regions in cf-mRNA during BM ablation and transplant. Day of blood collection with respect to transplant is indicated in the X axis. Dominant Ig transcripts, shown in solid blue and red lines, decrease after Melphalan-mediated BM ablation. (See also Fig S2).

To test whether cf-mRNA profiling can be used to monitor the levels of the malignant Ig clone, we sequenced the cf-mRNA from plasma of these patients every day for two weeks after chemotherapy and transplant. While Patient 1 showed no apparent reduction of the malignant clone after therapy, for Patient 2, decreasing levels of the predominant Ig variants were detected in cf-mRNA after Melphalan-induced apoptosis of plasma cells (Fig 2B-D and Fig S2A-C). By day 10, the immune profile was no longer dominated by clonal Ig combinations, indicating successful therapy and BM reconstitution (Fig 2B-D). In contrast, RNA-Seq performed on the matching buffy coat samples throughout the study showed very limited information regarding the malignant Ig transcripts (Fig2C, Fig S2A-C), revealing the potential of cf-mRNA to non-invasively capture BM activity over time.

### cf-mRNA captures hematopoietic lineage transcriptional activity during BM ablation and reconstitution

To gain further insights into the ability of circulating mRNA to reveal BM transcriptional activity, we followed the BM ablation and reconstitution dynamics after autologous transplants in cf-mRNA, using the prototypical MM Patient 2. Additionally, we investigated acute myeloid leukemia (AML) patients who underwent submyeloablative treatment followed by allogeneic transplant (see Materials and methods, AML Patients 1 and 2 were monitored for 8 weeks, Patient 3 was discharged 2 weeks after transplant). Unsupervised clustering of transcripts detected in cf-mRNA of MM and AML patients identified temporal patterns of expression for several groups of genes (Fig 3A, B). Both Gene Ontology enrichment analysis and RNA-seq data from Blueprint Consortium indicated that many of the identified components correspond to specific hematopoietic lineages (Fig 3A, B). Therefore, we examined in detail the dynamics of hematopoietic lineage-specific transcripts (i.e. erythrocytes, megakaryocytes, neutrophils) in circulation during BM ablation and reconstitution.

**Figure 3.**
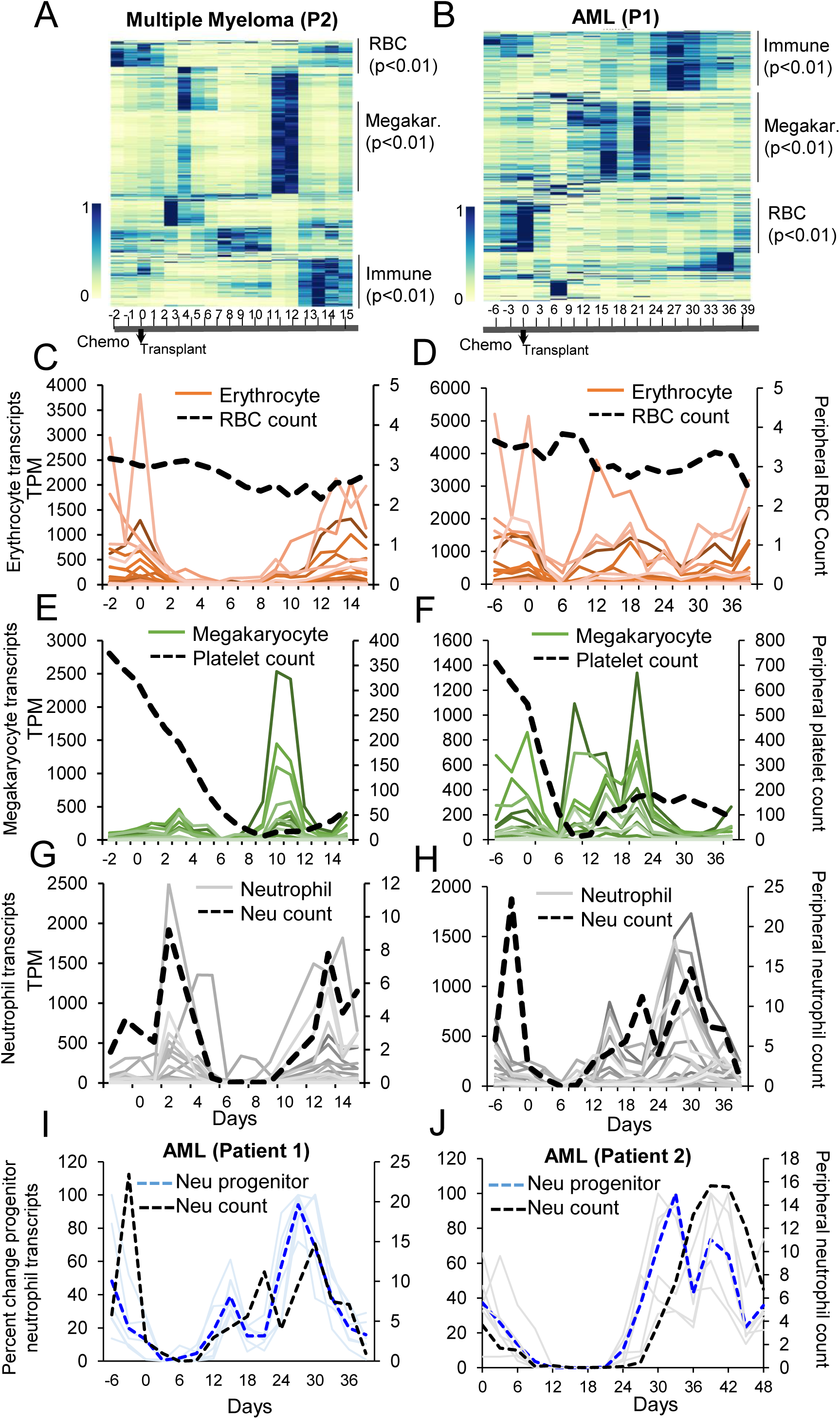
cf-mRNA reflects the transcriptional activity of hematopoietic lineages during BM ablation and reconstitution in cancer patients. A-B. Heat map of time-varying transcripts identified by cf-mRNA-Seq on multiple myeloma (MM) (A) and acute myeloid leukemia (AML) (B) patients undergoing BM ablation followed by autologous or allogenic stem cell transplant respectively (at day 0). Each column represents a time point with respect to the time of transplant, indicated in the bottom. Each row represent a gene. Enriched gene ontology terms for each cluster of trasncripts are indicated (adjusted p value). C-H. Time course of the levels of erythrocyte (red, C, D), megakaryocyte (green, E, F) and neutrophil (, gray, G, H) specific transcripts in MM (C, E, G) and AML (D, E, H) patients throughout the study. Transcript identity is provided in Table S2. Corresponding peripheral blood counts are plotted in the secondary axis and represented with a black dotted line (RBC count, millions per mL (C, D), platelet count, thousands per mL (E, F) and neutrophil count, thousands per mL (G, H). Day of blood collection with respect to transplant is indicated in the X axis. I-J. Relative variation of progenitor neutrophil transcripts in AML patients 1 (I) and 2 (J) throughout the study. Average percent change for these transcripts is represented with a dashed blue lane. Dashed black line shows neutrophil counts in blood. In both patients, during BM reconstitution progenitor neutrophil transcripts recovery in plasma precedes neutrophil count. (See also Fig S3, S4 and S5).

First, to clarify the relationship between erythrocyte circulating transcripts and RBCs, we examined the levels of erythrocyte lineage-specific transcripts in plasma and RBC counts throughout the study. RBCs are the predominant cell type in circulation and are stable for ~120 days in the bloodstream ^21^. Indeed, very little variation in RBC numbers was noticed in MM and AML patients during the duration of these studies (Fig 3C-D, Fig S4A). In contrast, erythrocyte-specific transcripts in cf-mRNA were rapidly reduced after chemotherapy-mediated BM ablation in all patients, and recovered at later time points during BM reconstitution (Fig 3C-D, FigS3 A-B, Fig S4A). The dramatic discrepancy between RBC number and erythrocyte transcripts in cf-mRNA indicates that these transcripts do not derive from circulating mature RBCs. Therefore, erythrocyte transcripts derive from immature erythrocyte forms either in the BM or in circulation (reticulocytes). We performed RNA-Seq analysis of paired buffy coat samples of MM Patient 2 to gain further insights into the origin of these transcripts. The levels of erythrocyte specific genes in CC are reduced after chemotherapy, resembling the dynamics observed in cf-mRNA (Fig S3C), and indicate that reticulocytes are the source of most erythrocyte transcripts in whole blood. However, transcripts like GATA1, a key transcriptional regulator of erythrocyte development, are clearly detectable in cf-mRNA earlier than in buffy coat during BM reconstitution (FigS3C), suggesting their BM origin. While future experiments will be necessary to discriminate the precise contribution of each compartment, our data shows that erythrocyte transcripts derive from immature erythrocyte cells residing in the BM and reticulocytes rather than from the highly abundant mature RBC.

To test whether the discrepancies between CBC and lineage-specific transcripts in circulation extend to other hematopoietic cell types, we next compared the dynamics of platelet counts and megakaryocyte-specific transcripts. In MM Patient 2, a dramatic increase in the levels megakaryocyte-specific transcripts is detected in cf-mRNA by day 9-10 after transplant, prior to platelet count recovery, which occurs by day 12-13 (Fig 3E). Interestingly, RNA-Seq from matched buffy coat samples showed that megakaryocyte transcript levels in CC mimic the dynamic of platelet counts throughout the study (Fig S3C), and, unlike in cf-mRNA, no early recovery of megakaryocyte transcripts is detectable in CC during BM reconstitution. This disparity suggests that megakaryocyte transcripts detected in cf-mRNA during BM reconstitution are not derived from CC, but from the BM. Supporting this observation, in AML Patient 1 megakaryocyte transcripts in circulation decreased after BM ablation and recovered by day 9, clearly foreshadowing the increase in platelet counts occurring by 12-13 (Fig3F). Strikingly, no recovery of this lineage occurred in cf-mRNA of AML Patient 2 (Fig S4B). Follow-up BM biopsy confirmed lack of megakaryocyte development in this patient (Table S3), showing the specificity of the measured megakaryocyte signal. Thus, our data indicates that cf-mRNA reflects megakaryocyte transcriptional activity in the BM during its reconstitution.

Last, we examined the kinetics of neutrophil counts and specific transcripts in circulation of MM and AML patients during the therapy. In MM Patient 2, neutrophil counts showed two spikes, one right after transplant, likely due to the G-CSF treatment, which is followed by a rapid decrease due to BM ablation, and a second spike by day 12, indicating BM reconstitution (Fig 3G). This resembles the overall dynamics of neutrophil-specific genes in cf-mRNA and in buffy coat during the procedure (Fig 3G, Fig S3E). However, while neutrophil transcripts in buffy coat and cf-mRNA peak at a similar time to neutrophil counts during BM reconstitution, neutrophil precursor genes like CTSG increased about 2 days earlier in cf-mRNA, by day 8-9 after the stem cell transplant. Supporting this observation, the levels of progenitor neutrophil transcripts in plasma of all AML patients decreased after BM ablation, and increased in cf-mRNA during BM reconstitution approximately 5 days earlier than the neutrophil counts (Fig 3 H-J and Fig S4D). These data further support that progenitor neutrophil transcripts in circulation are not derived from CC, but rather reflect BM transcriptional activity of the granulocyte lineage, providing valuable information about transplant engraftment and BM reconstitution.

We also investigated an orthogonal approach to measure transplant engraftment using cf-mRNA from AML patients receiving allogeneic HSC transplants, in which genetic differences exist between host and donor cells. Using a reference data base of SNPs we identified host specific polymorphisms in progenitor-neutrophil transcripts before the transplant (i.e. ELANE, AZU1, and PRTN3). After transplantation, these transcripts are substituted by new genetic variants from donor cells (Fig 4A). Indeed, cf-mRNA profiling enabled monitoring of changes in these transcripts during therapeutic treatment of Patients 1 and 2 (Fig 4B-C). Combined analysis of all detected SNP from the host switching to a different genetic variant after transplant (i.e. from homozygous to heterozygous) indicates that multiple genetic differences can be identified in cf-mRNA to temporally monitor transplant engraftment (Fig 4D-E). Altogether, our data shows that cf-mRNA captures both genetic information and transcriptional activity from the BM, and enables monitoring of ablation of malignant cells, transplant engraftment and BM reconstitution from donor cells during cancer treatment.

**Figure 4.**
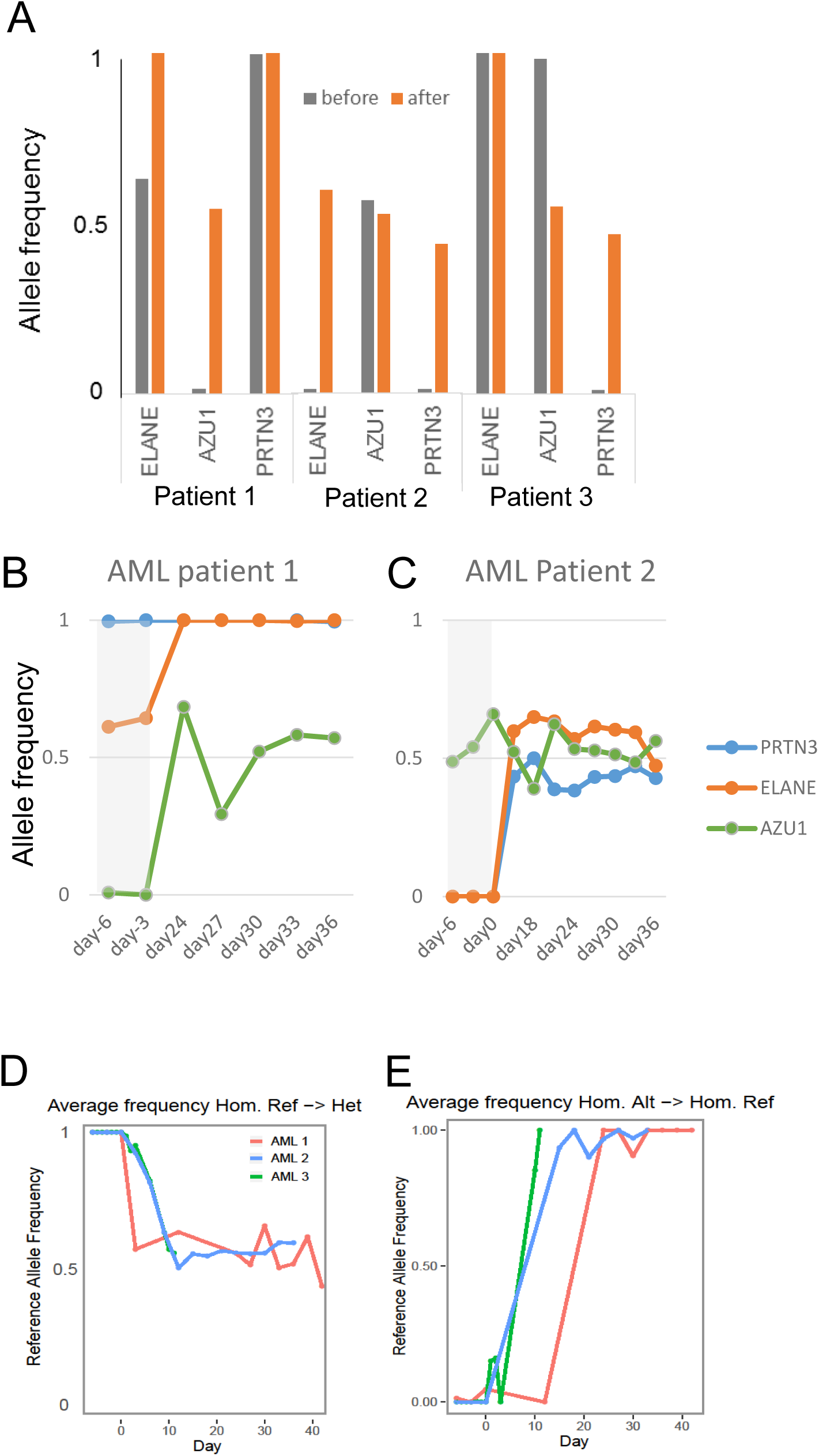
Monitoring of BM allotransplant engraftment in AML patients by genetic differences in cf-mRNA. A. Average frequency of reference allele of the SNPs detected in ELANE, AZU1 and PRTN3 neutrophil progenitor transcripts in cf-mRNA before (grey) and after (orange) allogeneic HSC transplantation in 3 AML patients, showing implantation of a new genetic profile after transplant. B-C. Frequency of reference allele of the SNPs detected in the same transcripts than in (A) for AML Patients 1 and 2. Day of blood collection with respect to the time of transplant is indicated in the X axis. D-E. Average reference allele frequency of all SNPs detected in the host cf-mRNA changing from reference homozygous to heterozygous (D) and from alternative homozygous to reference homozygous (E) after transplant. Day of blood collection is indicated in the X axis, transplant occurred at day 0.

### Lineage-specific transcriptional activity upon stimulation with growth factors is reflected in cf-mRNA

To evaluate the potential of cf-mRNA to monitor the activity of specific BM lineages after stimulation with growth factors, we obtained plasma from 9 patients with varying degrees of chronic kidney failure on chronic maintenance erythropoietin (EPO) therapy. EPO is a peptide hormone that specifically increases the rate of maturation and proliferation of erythrocytes in the BM ^22,23^. Samples were obtained prior to administration of EPO (day 0), and at several time points up to 30 days after treatment. Serum free hemoglobin and RBC number show minor transient changes during the duration of the study (data not shown). Unlike RBC counts, average levels of erythrocyte transcripts across 9 patients in cf-mRNA increased shortly after EPO treatment (Fig 5A). The levels of erythrocyte transcripts continued to increase during the initial days after treatment compared to untreated control individuals (Fig 5A, B). Indeed, key erythropoietic developmental transcripts involved in heme biosynthesis (i.e. ALAS2, HBB, HBA2) were induced in nearly all patients (8 out of 9 patients) (Fig S5A). Further, we identified 364 dysregulated genes in plasma by day 4 after treatment with EPO (p<0.05). Analysis using IPA (https://www.qiagenbioinformatics.com/products/ingenuitypathway-analysis) showed “Heme biosynthesis II” as the top enriched pathway for these transcripts (p<1.4e-9), supporting the transcriptional induction of this cell lineage. 30 days after EPO treatment, erythrocyte transcripts return to basal expression levels in these patients (Fig 5B and FigS5). Thus, our longitudinal studies indicate that cf-mRNA levels reflect specific transient stimulation of the erythroid lineage.

**Figure 5.**
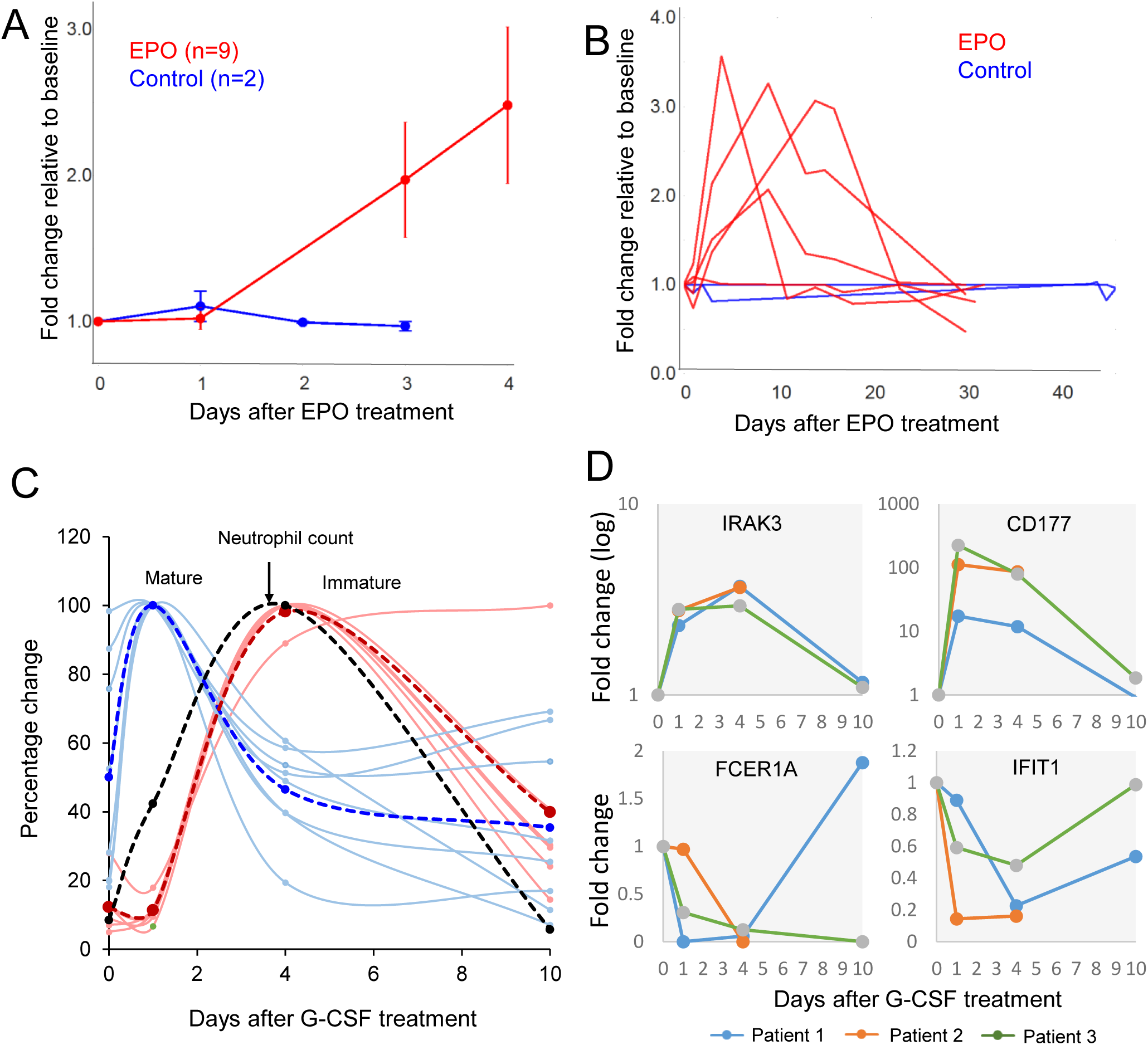
cf-mRNA captures the transcriptional activity of hematopoietic lineages upon stimulation. A. Blood was obtained from 9 patients before (day 0) and after (day 3, 4) being treated with a single EPO dose. Gene expression patterns in cf-mRNA were analyzed using RNA-Seq. Day 0 (before EPO treatment) was used as reference for each Patient, and changes in the levels of erythrocyte-specific transcripts after EPO treatment calculated. Average fold change of erythrocyte transcripts in all 9 patients subjected to EPO treatment (red) and 2 untreated controls (blue) are shown. B. Time course analysis of erythrocyte transcripts over a 30 day period in EPO treated patients (Red) Each line represents a patient, and shows average fold change of erythrocyte transcripts over time after a single EPO dosing administered at day 0, which is used as reference. Blue lines show fluctuations of the same transcripts in untreated healthy controls. See also Figure S6. C. Blood was obtained from 3 healthy patients treated with G-CSF (before treatment (day 0), and 1, 4 and 10 days after treatment). Changes in circulating transcriptome were analyzed by RNA-seq in plasma. Relative changes of immature (red) and mature (blue) neutrophil specific transcripts throughout the study are shown for a representative patient treated with G-CSF. Dashed red and blue lines indicate the average for each group of transcripts. Relative changes in neutrophil counts are shown in black. D. Time course of indicated G-CSF responsive genes measured by cf-mRNA-Seq. Plots show fold change over time relative to day 0. Time points are connected by lines, each line represent a patient. (See also Fig S6 and S7).

As another approach to study *in vivo* the effects in cf-mRNA of perturbation of a cell lineage, we collected samples from 3 healthy patients that received G-CSF treatment (granulocyte colony stimulating factor), a well-stablished pro-survival factor for neutrophilic granulocytes. Blood was drawn before the treatment and at 1, 4 and 10 days after G-CSF stimulation (the 10 day time point and CBC could only be obtained for 2 patients). As expected, neutrophil count increased after G-CSF treatment, peaking at day 4, and returned to basal levels by day 10 (Fig 5C). Neutrophil specific transcripts in cf-mRNA showed a bimodal increase after G-CSF treatment for all patients (Fig 5C and Fig S5 B,C). Neutrophil progenitor-specific transcripts increased in cf-mRNA coinciding with the peak in neutrophil counts as a consequence of G-CSF-mediated mobilization of granulocytes from the BM into circulation (Fig 5C, Fig S5B). However, mature neutrophil transcripts rapidly increase in cf-mRNA 1 day after the treatment, foreshadowing the peak of neutrophil counts (Fig 5C, Fig S5C). This suggests a direct and transient transcriptional response of neutrophils to G-CSF. Indeed, transcripts previously reported both *in vivo* and *in vitro* to increase (e.g., IRAK3) or decrease (e.g., IFIT1) in neutrophils in response to G-CSF, followed the expected transcriptional response ^24^(Fig 5D). Altogether, our results indicate that cf-mRNA reflects cell type-specific transcriptional responses to stimulation.

## Discussion

The growing interest to identify non-invasive alternatives to standard tissue biopsies has generated enormous attention for “liquid biopsies” over the last decade. Initial advances in cf-DNA technology have paved a way for the development of clinically applicable cf-NA-based biomarkers ^25-27^. cf-DNA offers potential advantages compared to invasive tissue biopsies; however, cf-DNA analyses largely rely on mutations, polymorphisms, or structural variation, preventing its use in disease and physiological scenarios not associated with genetic differences. To partially circumvent these limitations, cf-DNA methylation analyses have recently been used as a surrogate of tissue-specific gene expression, but further work will be needed to validate this approach ^28^. In contrast, the cf-mRNA transcriptome provides direct access to both genetic information as well as information pertaining to the tissue of origin and its physiology. For instance, we have shown that genetic alterations in cf-mRNA provide valuable information for monitoring allografts, and similar approaches have shown their value in diagnosing fetal chromosomal abnormalities ^29^. Given that several studies have identified tumor derived transcripts in circulation ^14^, the genetic information captured by cf-mRNA is of special interest in cancer diagnosis and monitoring. Indeed, we were able to detect mutations in cf-mRNA identified during routine molecular testing in AML patients (data not shown). In addition, cf-mRNA provides tissue-specific transcripts that reveal functional information about the tissue of origin. We show that cf-mRNA captures transcripts that reveal BM physiology in both healthy subjects and cancer patients. Similarly, previous studies have reported transcripts in circulation encoding functional information of the liver, brain, immune system or fetal development ^7,30,31^. Therefore, cf-mRNA has the capability of integrating functional and genetic information of tissues, highlighting its unique potential as a non-invasive biomarker.

Another key aspect of non-invasive approaches is that by eliminating the need for surgical tissue acquisition they enable repeated assessment of a patient’s disease state over time. This could be of great significance in several clinical settings, such as monitoring of treatment in cancer patients, where biopsy of affected tissue remains gold standard. In this regard, our longitudinal cf-mRNA profiling data shows that circulating transcripts capture snapshots of gene expression of tissues such as BM. This allows non-invasive temporal delineation of BM ablation efficiency, early detection of transplant engraftment and monitoring BM reconstitution. For example, in MM patients, cf-mRNA profiling integrates temporal measurement of clonal Ig transcripts generated by malignant plasma cells in the BM, with detailed BM-lineage transcriptional activity and establishment of a new immune profile. The comprehensive picture revealed by cf-mRNA profiling provides additional clinically relevant information compared to other non-invasive tests commonly used in this malignancy, such as clonal antibody detection in serum of MM patients. Indeed, given the challenging and subjective quantification and characterization of these antibodies, BM biopsies remain as common practice in the therapy management of MM patients ^32^. In addition, unlike antibody detection, cf-mRNA profiling has the potential for early identification of suboptimal BM reconstitution, as shown by the lack of development of megakaryocyte lineage in AML Patient 2. Our data indicate that cf-mRNA profiling could be translated into the clinical setting to improve the therapeutic management of cancer patients, alleviating the need for invasive BM biopsies.

Finally, understanding the mechanisms underlying the presence of mRNA transcripts in circulation is essential to interpret their clinical value. For example, cf-DNA has been shown to originate primarily from dying cells ^12^; therefore, the use of this liquid biopsy strongly relies on scenarios associated with cell death. Strikingly, we showed that melphalan-induced apoptosis did not significantly increase the levels of cf-mRNA. In contrast, a large increase of transcripts in circulation was observed during BM reconstitution and upon stimulation with well-known pro-survival and antiapoptotic growth factors. These data indicate that changes in cf-mRNA levels are heavily influenced by active transcriptional changes in living cells during maturation, proliferation and response to stimuli, without requiring cell death. Supporting this observation, *in vitro* studies have shown that extracellular mRNA levels and composition change upon cellular stimulation ^33,34^. Additionally, our clinical longitudinal studies demonstrate that the circulating transcriptome is a dynamic compartment, which allows constant measurement of tissue function over time. This is in contrast to cf-DNA methylation and mutation events, which are less dynamic and provide limited information on tissue homeostasis. Therefore, cf-mRNA profiling provides broader molecular information compared to other non-invasive biomarkers and constitutes a unique non-invasive approach to examine tissue function in scenarios such as monitoring of diseases and drug response in patients.

## Acknowledgements

We thank the San Diego Blood Bank for providing samples.

## Author contributions

Conceptualization MN, Methodology AI, NSS, YZ; Investigation AI, NSS, VH, AA, APK, LG, JRP and IP; Writing, AI; Resources; JA, TSN, MN, JM, DS, SRQ; Funding acquisition TSN, MN, JA and SRQ. Visualization, AI, ST and JZ; Formal analysis: YZ, JZ. Project administration, AI, JA; Supervision: MN

