## Supplemental figures and figure legends for "Non-invasive characterization of human bone marrow by cell free messenger-RNA reveals response to growth factor stimulation and hematopoietic reconstitution after transplantation"

**Supplemental Materials:**

**-Supp. Fig. Legends**

**-Supplemental Tables**

**Figure S1. cf-mRNA transcriptome is enriched in bone marrow transcripts compared to circulating cell transcriptome.**

A. Schematic of whole blood, plasma and buffy coat isolation.

B-C. Scatter plots comparing the levels in peripheral blood (X axis) and cf-mRNA (Y axis) of neutrophil-specific and T-cell-specific transcripts. Neutrophil progenitor transcripts are shown in red, mature transcripts in blue. Both x-axis and y-axis show TPM in $\log_2$ scale.

D-E. Box-plots comparing the normalized levels (TPM) of the indicated hematopoietic progenitor transcripts measured by RNA-Seq in paired buffy coat and cf-mRNA samples (n=5; p-value, t-test)

F. Levels of BM-specific (red dots) and peripheral blood-specific genes (blue dots) were compared in matching plasma and peripheral blood of 3 individuals. Average fold change (plasma/whole blood) of these transcripts is shown. P value, t test.

**Figure S2. cf-mRNA contains Ig transcripts derived from plasma cells in the BM of Multiple Myeloma patients.**

A-C. Levels of Ig transcripts measured by RNA-Seq in plasma and buffy coat of a MM patient undergoing BM ablation (starting day -2) and autologous stem cell transplantation (day 0). Bar graphs show the normalized levels (TPM) of Ig heavy chain constant region transcripts (A), light chain constant region transcripts (B) and lambda light chain variable region transcripts (c) detected during the study. Day of blood collection with respect to the time of transplant is indicated in the X axis. Ig transcripts IGHG1 and IGKC dominate the plasma sample, matching the results obtained by molecular testing performed in BM biopsy of this patient (Table S3).

A. Fraction of Ig heavy and light variable chain transcripts over time in cf-mRNA of MM Patient 1. Dominant transcripts are shown in solid blue and red lines. Time with respect to transplant day is shown.

**Figure S3. Monitoring BM transcriptional activity by cf-mRNA profiling in a Multiple Myeloma patient during BM ablation and transplant**

A-B. Time course of red blood cell counts (RBC, dashed black line) and hemoglobin transcripts (grey) in multiple myeloma Patient 2 during chemotherapy and BM reconstitution (see also Fig 3). Day of blood collection with respect to the time of transplant is indicated in the X axis.

C-F. RNA-Seq was performed in cf-mRNA (blue) and matching buffy coat samples (red). Graphs show the fold change relative to baseline of key erythrocyte (C) and megakaryocyte transcripts (D), as well as mature neutrophil (E) and immature neutrophil-specific transcripts (F) in both specimens. In all panels, black lines represent the relative changes in corresponding circulating cell blood counts: RBC counts (C), platelet counts (D) and neutrophil counts (E, F). Day of blood collection with respect to the time of transplant is indicated in the X axis.

**Figure S4. Monitoring transcriptional activity of BM hematopoietic lineages by cf-mRNA in Acute Myeloid Leukemia (AML) patients undergoing BM ablation and transplant.**

A-C. Time course of normalized levels (TPM) of erythrocyte (A), megakaryocyte (B) and neutrophil (C) specific transcripts in AML Patient 2 (grey lines). Corresponding peripheral blood counts are plotted in the secondary axis of each graph and represented with a black dotted line (RBC count (A), platelet count (B) and neutrophil count (C). Day of blood collection with respect to the time of transplant (day 0) is indicated in the X axis.

D.Time course of mature (dark blue) and immature neutrophil components (light blue) in AML patients. Neutrophil count is shown in red. Immature transcripts are detected in cf-mRNA days before neutrophil count recovers. Day of blood collection with respect to the time of transplant is indicated in the X axis.

**Figure S5. Induction of lineage specific-genes in cf-mRNA by growth factors after EPO treatment**

A. Fold change over time of key erythrocyte developmental genes (indicated) in EPO treated patients relative to baseline. The general trends show elevated levels of these transcripts after EPO treatment with a return to basal levels at later time points.

B-C. Fold change of immature (A) and mature (B) neutrophil specific transcripts in cf-mRNA of a patients after treatment with G-CSF. Day 0 (before treatment) is used as reference. Fold change of indicated transcripts is shown for 3 patients, represented in different colors. Time points across each Patient are connected by lines. Day of blood collection with respect to the time of treatment is indicated in the X axis.

Table S1. Number of transcripts detected in cf-mRNA of healthy donors (n=24).

| | >40% of the samples | >60% of the samples | >80% of the samples |
|---|---|---|---|
| TPM > 1 | 12341 | 11393 | 10313 |
| TPM > 5 | 9414 | 8485 | 7334 |

TPM (Transcripts Per Million)

Table S2. List of indicated hematopoietic lineage-specific transcripts.

| Erythrocyte | Megakaryocyte | T-cells | T-cells | T-cells | Neutrophil | Immature neutrophil | Mature neutrophil |
|---|---|---|---|---|---|---|---|
| SLC4A1 | ITGA2B | PDZD4 | TRGV10 | TRAV23DV6 | PGLYRP1 | ELANE | S100A12 |
| TF | RAB27B | TBX21 | TRGV4 | TRAV26-1 | LTF | PRTN3 | KRT23 |
| AVP | GUCY1B3 | CHRNA3 | TRBV6-1 | TRAV41 | ATP2C2 | AZU1 | FCGR3B |
| RUNDC3A | GP6 | SIRPG | TRBV9 | DBH-AS1 | VNN3 | CTSG | PI3 |
| SOX6 | HGD | PITPNM2 | TRBV6-5 | AC011893.3 | CRISP3 | RNASE | STEAP4 |
| TSPO2 | PF4 | GZMH | TRBV5-6 | RP11-73O6.3 | CTSG | PGLYRP1 | PROk2 |
| HBZ | CLEC1B | GZMB | TRBV4-2 | TRBV10-2 | OLFM4 | MMP8 | CXCR1 |
| TMCC2 | CMTM5 | GZMK | TRBV20-1 | TRBV5-4 | KRT23 | | CXCR2 |
| SELENBP1 | GP9 | GNLY | TRBC1 | RP11-144L1.4 | MMP8 | | CD177 |
| ALAS2 | SELP | CD2 | TRBV27 | LINC00987 | ARG1 | | KCNJ15 |
| EPB42 | DNM3 | CD160 | TRAV2 | TRBV30 | EPX | | ALPL |
| GYPA | LY6G6F | ELOVL4 | TRAV3 | TRBV3-1 | PI3 | | |
| C17orf99 | LY6G6D | EPHX2 | TRAV4 | TRBV11-2 | CRISP2 | | |
| HBA2 | XXbac-BPG32J3.19 | SARDH | TRAV10 | A2M-AS1 | STEAP4 | | |
| RHCE | RP11-879F14.2 | KLRC1 | TRAV12-2 | LINC01550 | LCN2 | | |
| HBG2 | | FGFBP2 | TRAV13-2 | RP11-291B21.2 | PRG3 | | |
| TRIM10 | | ARL5C | TRAV14DV4 | TRAV1-2 | KCNJ15 | | |
| HBA1 | | RORC | TRAV12-3 | RP11-204N11.1 | ALPL | | |
| HBM | | GZMA | TRAV17 | RP11-158G18.1 | FCGR3B | | |
| HBG1 | | SCML4 | TRAV19 | RP11-415F23.3 | S100A12 | | |
| UCA1 | | EPHA1 | TRAV20 | TRBV15 | PROK2 | | |
| GYPB | | KLRF1 | TRAV21 | TRBV12-4 | CXCR1 | | |
| CTD-3154N5.2 | | PPP1R1C | DTHD1 | CXCR6 | CAMP | | |
| AC104389.1 | | CD8A | KLRC2 | THEMIS | RNASE3 | | |
| | | PPP2R2B | RP11-415F23.4 | LRRN3 | CEACAM3 | | |
| | | TRAT1 | RP11-104L21.3 | CCR9 | AZU1 | | |
| | | CTLA4 | TRBV12-3 | PRF1 | ABCA13 | | |
| | | MAL | TRBV10-3 | FCRL6 | CXCR2 | | |
| | | CD8B | TRBV13 | TIGIT | CTD-3088G3.8 | | |
| | | | | ADARB2 | PRTN3 | | |
| | | | | | ELANE | | |
| | | | | | CD177 | | |
| | | | | | LINC00671 | | |
| | | | | | ORM2 | | |
| | | | | | ORM1 | | |
| | | | | | HP | | |
| | | | | | RP11-678G14.4 | | |

Table S3: Patient characteristics

**Multiple myeloma patient characteristics**

| Patient | 1 | 2 | 3 |
|---|---|---|---|
| Age | 75 | 52 | 67 |
| Sex | Male | Male | Female |
| Diagnosis | IgA lambda | IgG Kappa | IgA Kappa |
| Peak relevant Ig prior to treatment | 0.6 g/dl | 5.6 g/dl | 1.4 g/dl gamma |
| Plasma cells at time of transplant | 13% | 1% | <1% |
| Prior treatment | Radiation, VRD | Radiation, VRD | VRD |
| Plasma cells after treatment | N/A | <0.5% | <1% |
| Relevant Ig after transplant | 0.16 g/dl | 0.8 g/dl | 0.038 g/dl |

VRD-bortezomib, lenalidomide, dexamethasone

After transplant: evaluated 60 d post-procedure

**: AML patient characteristics**

| Patient | 1 | 2 | 3 |
|---|---|---|---|
| Age | 68 | 66 | 66 |
| Sex | Female | Male | Male |
| Bone marrow blast (%) | 16 | 3 | 50 |
| Prior Therapy | Yes* | No | No |
| Additional information | | ** | |

*diffuse large B-cell lymphoma

** BM biopsy revealed lack of megakaryocyte development in Patient 2

**: EPO patient characteristics**

| Patient No | 1 | 2 | 3 | 4 | 5 | 6 | 7 | 8 | 9 |
|---|---|---|---|---|---|---|---|---|---|
| Age | 84 | 67 | 82 | 91 | 73 | 78 | 80 | 74 | 80 |
| Chronic kidney disease stage | 4 | PD | 4 | 4 | 3 | 4 | 3 | 5 | 3 |
| EPO agent | Aranesp | Procrit | Aranesp | Procrit | Aranesp | Aranesp | Aranesp | Procrit | Aranesp |
| Creatinine concentration (md/dL) | 1.8 | 4.1 | 2.7 | 2.3 | 1.3 | 2.4 | 1.1 | 4.5 | 1.5 |

PD- Peritoneal Dialysis

**: G-CSF patient characteristics**

| Patient No | 1 | 2 | 3 |
|---|---|---|---|
| Age | 56 | 34 | 24 |

**Figure S1**

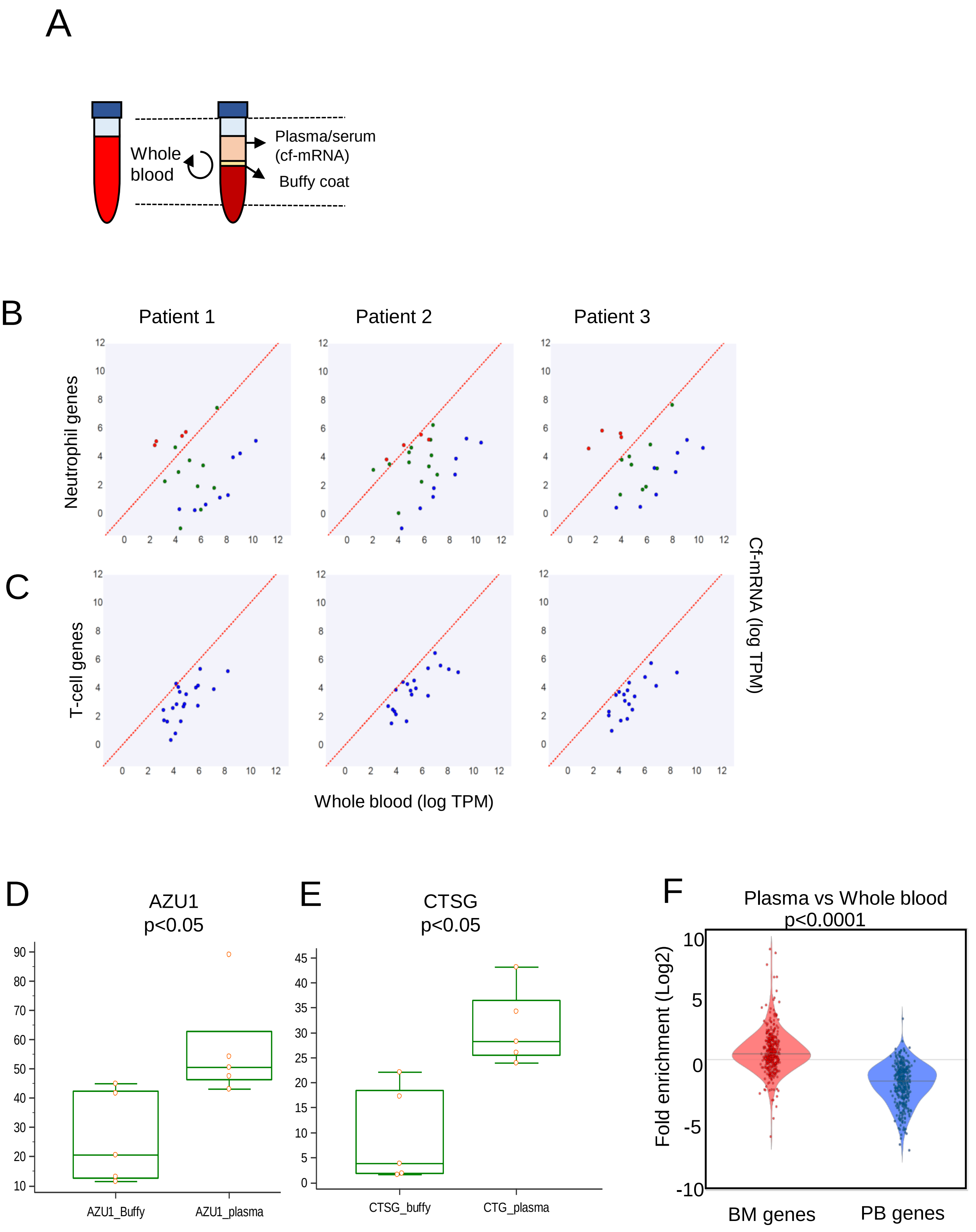
A
Whole blood
Plasma/serum
(cf-mRNA)
Buffy coat
B
Patient 1
Patient 2
Patient 3
Neutrophil genes
C
T-cell genes
Cf-mRNA (log TPM)
Whole blood (log TPM)
D
AZU1
p<0.05
AZU1_Buffy
AZU1_plasma
E
CTSG
p<0.05
CTSG_buffy
CTG_plasma
F
Plasma vs Whole blood
p<0.0001
Fold enrichment (Log2)
BM genes
PB genes

### Figure S2

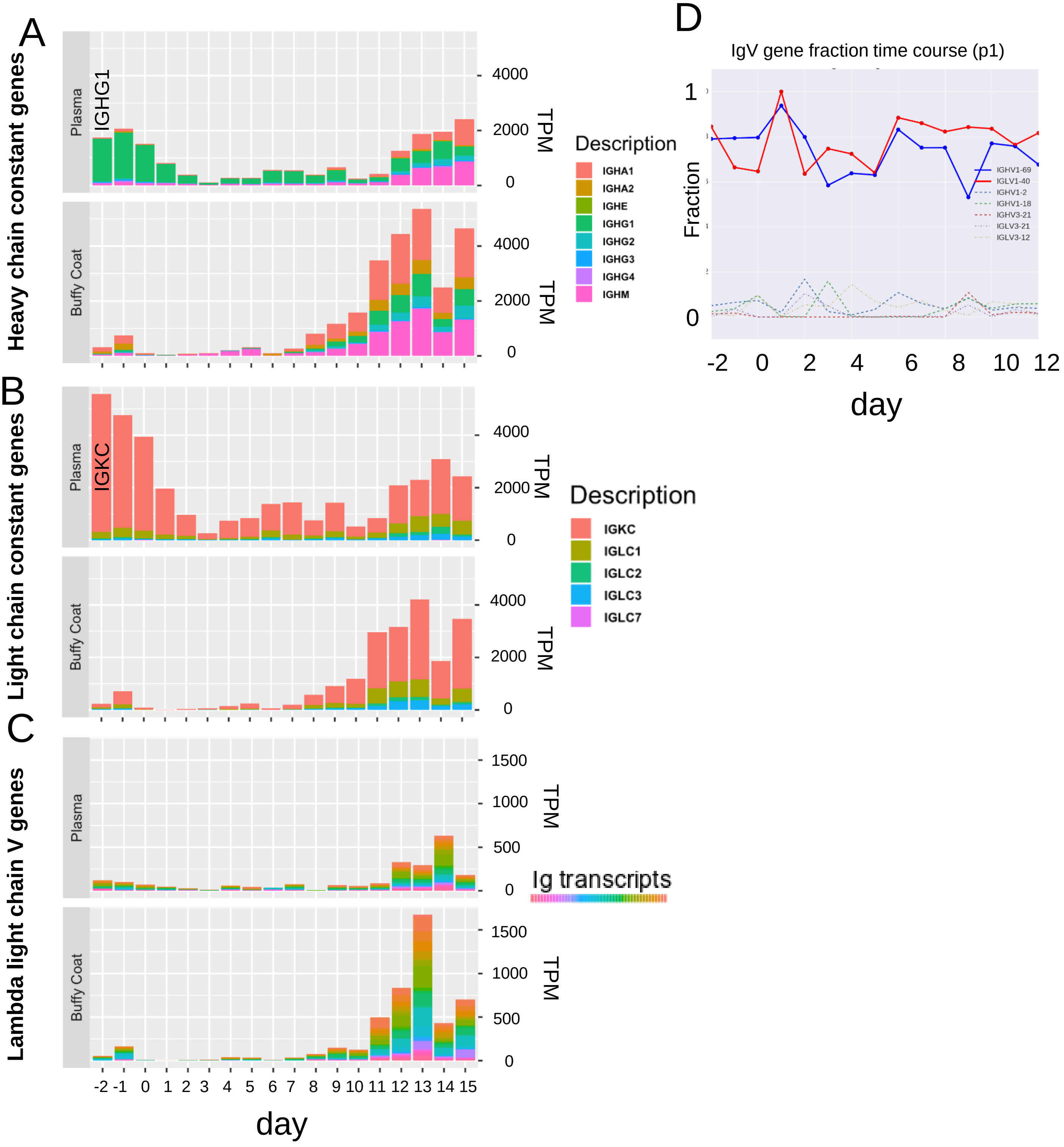

A
Heavy chain constant genes
Plasma
IGHG1
Buffy Coat
TPM
4000
2000
0
Description
IGHA1
IGHA2
IGHE
IGHG1
IGHG2
IGHG3
IGHG4
IGHM
B
Light chain constant genes
IGKC
IGLC1
IGLC2
IGLC3
IGLC7
C
Lambda light chain V genes
1500
1000
500
Ig transcripts
-2 -1 0 1 2 3 4 5 6 7 8 9 10 11 12 13 14 15
day
D
IgV gene fraction time course (p1)
Fraction
1
0
-2 0 2 4 6 8 10 12
IGHV1-69
IGLV1-40
IGHV1-2
IGHV1-18
IGHV3-21
IGLV3-21
IGLV3-12

### Figure S3

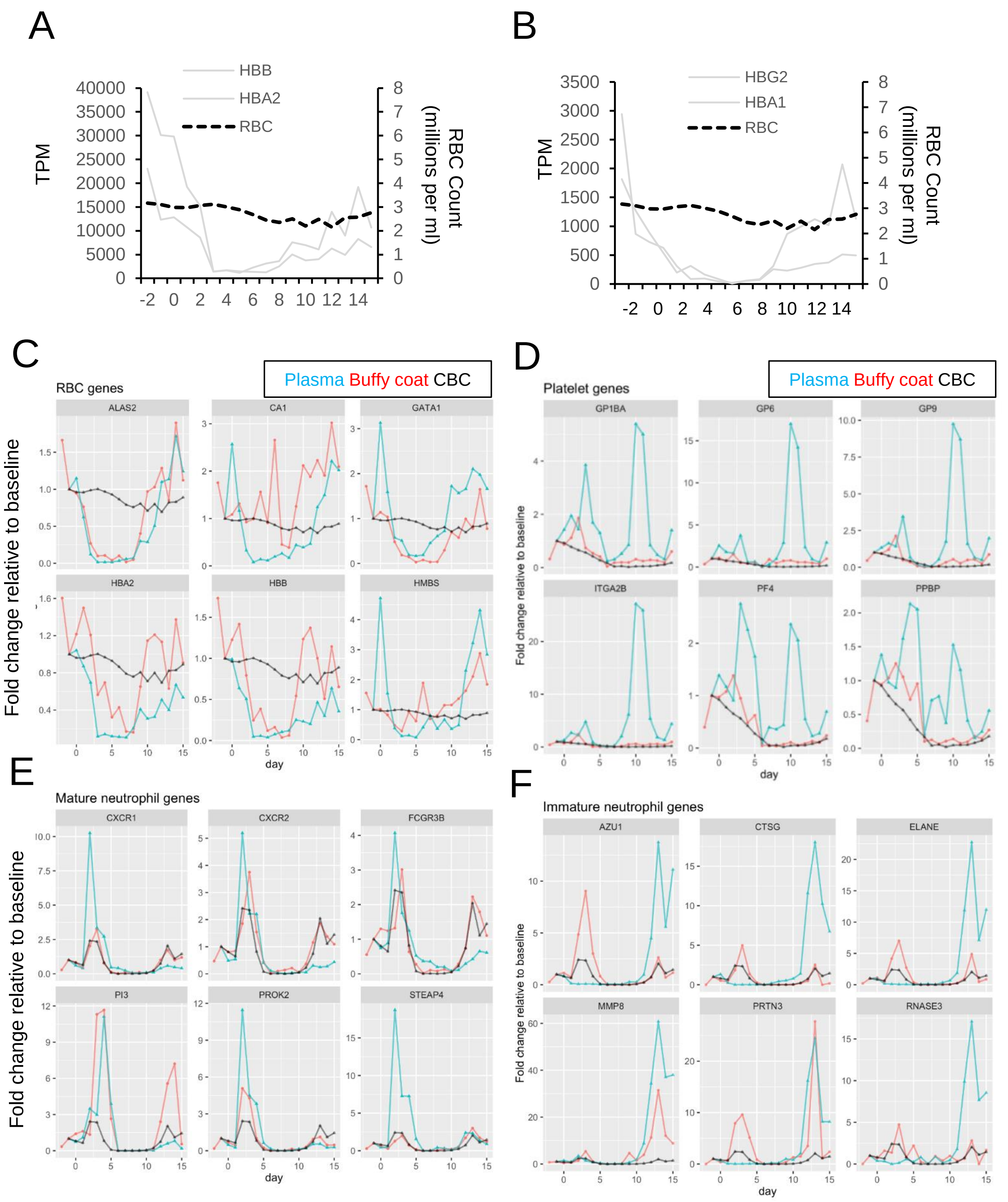
A
HBB
HBA2
RBC
TPM
RBC Count (millions per ml)
B
HBG2
HBA1
RBC
TPM
RBC Count (millions per ml)
C
Plasma Buffy coat CBC
RBC genes
ALAS2
CA1
GATA1
HBA2
HBB
HMBS
Fold change relative to baseline
day
D
Plasma Buffy coat CBC
Platelet genes
GP1BA
GP6
GP9
ITGA2B
PF4
PPBP
Fold change relative to baseline
day
E
Mature neutrophil genes
CXCR1
CXCR2
FCGR3B
PI3
PROK2
STEAP4
Fold change relative to baseline
day
F
Immature neutrophil genes
AZU1
CTSG
ELANE
MMP8
PRTN3
RNASE3
Fold change relative to baseline
day

### Figure S4

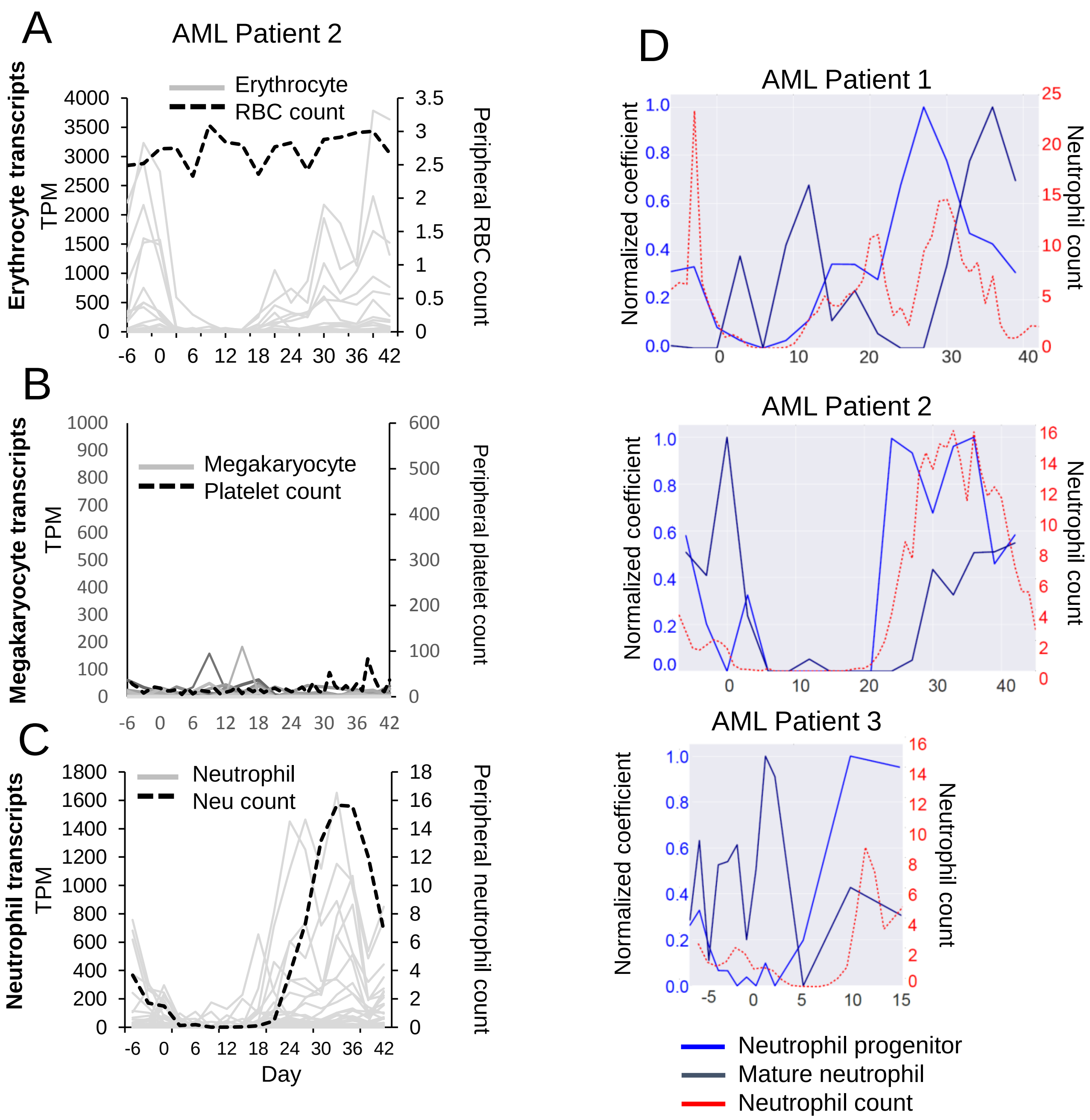
A
AML Patient 2
Erythrocyte
RBC count
Erythrocyte transcripts
TPM
Peripheral RBC count
B
Megakaryocyte
Platelet count
Megakaryocyte transcripts
TPM
Peripheral platelet count
C
Neutrophil
Neu count
Neutrophil transcripts
TPM
Peripheral neutrophil count
Day
D
AML Patient 1
AML Patient 2
AML Patient 3
Normalized coefficient
Neutrophil count
Neutrophil progenitor
Mature neutrophil
Neutrophil count

### Figure S5

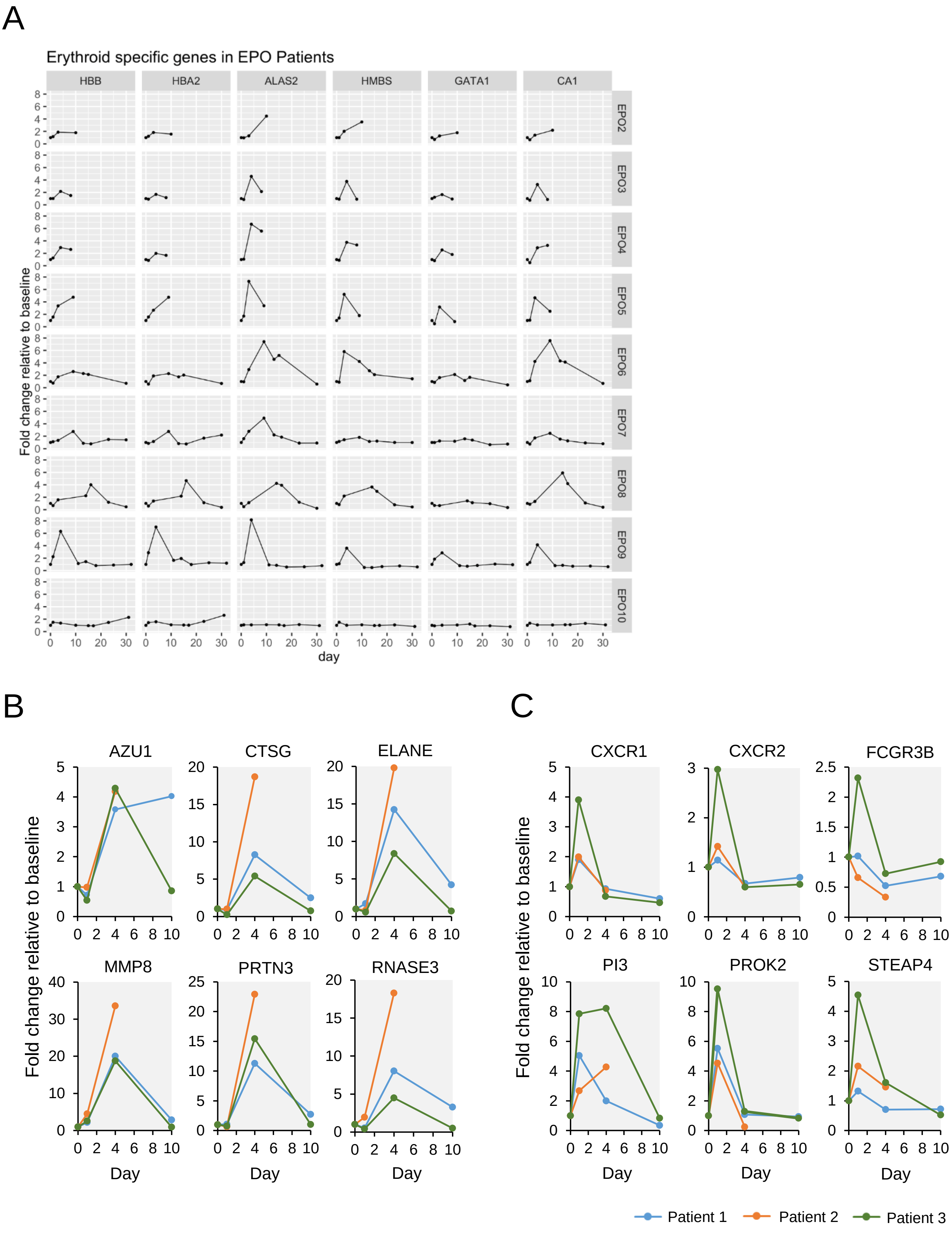
A
Erythroid specific genes in EPO Patients
HBB
HBA2
ALAS2
HMBS
GATA1
CA1
EPO2
EPO3
EPO4
EPO5
EPO6
EPO7
EPO8
EPO9
EPO10
Fold change relative to baseline
day
B
AZU1
CTSG
ELANE
MMP8
PRTN3
RNASE3
Fold change relative to baseline
Day
C
CXCR1
CXCR2
FCGR3B
PI3
PROK2
STEAP4
Fold change relative to baseline
Day
Patient 1
Patient 2
Patient 3
